# GIN-CRC-Pareto: A graph-based Pareto-optimal multi-task learning framework to identify miRNA-target interactions in colorectal cancer

**DOI:** 10.1101/2025.08.10.669528

**Authors:** Lin Li, Qiang Yang, Lu Li, Hongru Zhao, Jie Xu, Mingyi Xie, Rui Yin

## Abstract

Colorectal cancer (CRC) ranks as the third highest incidence among malignancies in humans and the second most common cause of cancer-related mortality in the United States. Accumulating evidence has established microRNAs (miRNAs) as critical regulators of cancer development and therapeutic response. Understanding miRNA-mRNA interactions is critical for elucidating the molecular mechanisms driving CRC and other malignancies. In this study, we proposed GIN-CRC-Pareto, a graph-based, Pareto-optimal multi-task learning framework that simultaneously predicts miRNA-mRNA binding pairs, identifies seed match pairings, and classifies seed match subtypes. By leveraging the power of graph neural networks and Pareto-optimal gradient balancing strategy, GIN-CRC-Pareto dynamically adjusted the task weights during training to optimize each task without compromising the others. Experimental results demonstrated that our framework consistently outperforms traditional deep learning models and existing state-of-the-art tools across multiple evaluation metrics, with 0.909 in accuracy, 0.909 in precision and 0.968 in AUC in the miRNA-mRNA binding pairs prediction task. Additionally, we further validated the generalizability of the framework in combination with transfer learning techniques to identify miRNA-target interactions across other cancers. These findings highlight the effectiveness of the proposed framework to comprehensively identify the miRNA-target interactions in CRC, with the potential to serve as a scalable and generalizable tool across diverse cancer types, ultimately facilitating the development of miRNA-based therapeutics for cancer treatment.

## Introduction

Colorectal cancer (CRC) presents a significant global health burden as the third most common cancer and the second leading cause of cancer-related deaths. In 2024, there were an estimated 152,810 new cases of CRC in the United States, causing about 53,010 deaths [1]. Alarmingly, CRC incidence rates among younger adults (aged <55 years) have been increasing by 1%–2% annually since 2005 [2]. This trend is expected to worsen in the near future, driven by aging population and lifestyle changes, particularly in middle- and low-income countries [3]. Moreover, diagnoses have increasingly shifted toward more advanced stages. Since 2010, the proportion of CRC cases diagnosed at regional or distant stages rose from 52% in the mid-2000s to 60% in 2021 [2]. This increase highlights the challenge of detecting early symptoms, often leading to diagnoses only after significant disease progression. Given the growing prevalence of CRC, understanding its genetic and molecular basis is critical for developing effective strategies for early detection and treatment. MicroRNAs (miRNAs), known as small, single-stranded non-coding RNAs, have been consistently revealed to regulate gene expression and play crucial roles in numerous biological processes [4]. Dysregulation of miRNAs is related to many diseases, such as neurological disorders [5], retinal disorders [6], cancer [7], and cardiovascular diseases [8]. In the context of CRC, miRNAs contribute to tumorigenesis or tumor suppression by targeting specific mRNAs, making them potential biomarkers for screening and diagnostics [9,10]. For instance, a newly identified miRNA, i.e., miR-124, significantly reversed cancer characteristics (proliferation, stemness, migration, and invasion) in CRC cells by reducing STAT3 signaling activity and altering its downstream targets [11], indicating its potential tumor-suppressive effect. Other CRC-associated miRNAs include miR-708 and miR-148a-3p. It was reported that miR-708 suppresses CRC development via regulating ZEB1 in the AKT/mTOR pathway [12], while miR-148a-3p was demonstrated to target PD-L1 to prevent tumorigenesis [13].

The miRNA-mRNA binding pairs can be generally classified into two categories, that is, seed region and non-seed region interactions. The seed region interaction, known as canonical interaction, occurs when nucleotides 2-8 of the miRNA show complementarity to the 3’ untranslated region (3’UTR) of the target mRNA [14]. This type of interaction is typically associated with strong regulatory effects, leading to mRNA degradation or translational repression. It can be further categorized into several types based on the extent and position of the match within the seed region: 8-mers (positions 1–8 with an adenine at target position 1), 7-mer-m8 (positions 2–8), 7-mer-A1 (positions 1-7 with an adenine at target position 1), 6-mers (positions 2–7), and offset-6-mer (positions 3–8). The regulatory efficacy of these subtypes follows a hierarchical order: 8-mer > 7-mer-m8 > 7-mer-A1 > 6-mer > offset-6-mer [15]. In contrast, non-seed region interactions introduce additional complexity and flexibility in miRNA-mediated regulation. For instance, compensatory interactions feature imperfect seed pairing that is counterbalanced by strong 3’ region binding, enabling sustained regulatory function. These diverse interaction mechanisms underscore the dynamic nature of miRNA-mediated gene regulation, allowing for context-dependent and tissue-specific control of gene expression. Despite the existence of non-canonical binding, a substantial number of mRNAs contain sequences complementary to the miRNA seed region, reaffirming its central role in regulatory specificity [16]. However, computational predictions of non-seed interactions are more susceptible to false positives [17]. This hierarchy is also evident at the protein level, where the most strongly repressed proteins are typically linked to mRNAs harboring at least one 7-mer or 8-mer site in the 3′-UTR [18,19]. Therefore, accurately identifying the category and subtypes of seed match types is essential for deciphering the regulatory specificity of miRNA-mRNA interactions.

The advent of high-throughput experimental techniques, such as AGO-CLIP-seq (Argonaute cross-linking and immunoprecipitation sequencing) [20] and CLASH (Crosslinking, Ligation, and Sequencing of Hybrids) [21], can provide experimentally validated interactions between miRNAs and their targets. However, these methods are generally extremely laborious, time-consuming, and expensive [22,23]. Computational approaches have emerged as scalable alternatives for miRNA target prediction that can reduce experimental workload and enhance efficacy. For example, Wen et al. developed DeepMirTar, using a stacked denoised autoencoder to characterize miRNAs and their candidate target sites [24]. Pla et al. introduced miRAW [25], which applied deep artificial neural networks (ANN) for miRNA-target prediction. Yu et al. developed preMLI [26], integrating rna2vec, CNN, and BI-GRU models to identify miRNA– lncRNA binding pairs. More recently, Yin et al. illustrated the effectiveness of combining GNNs with next-generation sequencing data to uncover miRNA-mRNA binding interactions in CRC, emphasizing the power of graph-based methods to extract hidden relationships [27]. Despite these advances, most existing methods focused on identifying whether there occurs miRNA–target binding pairs without further elucidating more detailed binding categories. For instance, while seedmatchR [28] could predict the seed match subtypes of miRNA-target interactions (MTIs), the model was designed primarily for siRNA-mRNA interactions, and the analysis relied on statistical and computational string matching methods. Further, deep learning models have been applied to reveal the categories of MTIs through candidate target site prediction. Min et al. proposed Targetnet [29], which employed deep residual network (ResNet) with one-dimensional CNN to predict the miRNA candidate target site. Zhang et al, developed miTDS [30] that combined CNN and BiLSTM model with transformer-based semantic features to identify functional target sites. Nonetheless, limited attention has been given to the inductive bias between different tasks of binding site prediction. The knowledge learned from the binary miRNA-mRNA binding pairs classification task can be beneficial to the seed match subtype prediction task by helping the model identify generalizable patterns and features, such as sequence motifs and structural similarities. These shared features could serve as a foundation for more granular classification, acting as a regularization mechanism and enhancing model generalizability. This is particularly crucial for small but biologically significant seed match subtypes, such as 8-mer, which account for only 1.42% of miRNA-mRNA binding pairs in CRC. However, implementing a multi-task learning framework to balance these tasks is nontrivial. A common strategy involves using hyperparameters to control the relative importance of tasks during training, but this approach is only valid when the tasks are non-competitive, which is uncommon. The binary classification task may emphasize macro-level distinctions at the expense of nuanced patterns critical for subtype classification. This conflict potentially introduces biases or oversimplifications that challenge effective knowledge transfer between tasks.

To address the challenges in miRNA target prediction tasks, we proposed a graph-based Pareto-optimal multi-task learning framework named GIN-CRC-Pareto that extends beyond conventional binary miRNA-target binding pairs prediction. In multi-task optimization, a solution is considered Pareto-optimal when no task can be improved without degrading the performance of at least one other task [31]. We employed this principle by dynamically adjusting task weights during training, ensuring that improvements in one task do not come at the cost of others, promoting an equitable balance that aligns with the biological relevance of the tasks [32]. More specifically, our framework concurrently addressed three tasks: (1) predicting the presence of a miRNA-mRNA binding pair, (2) identifying whether the interaction involves a seed match pairing, and (3) classifying the specific seed match subtypes, with a focus on CRC. By integrating these tasks within a unified multi-task learning model, we aimed to capture both high-level interaction patterns and fine-grained biological insights essential for comprehensive miRNA target prediction. This framework leveraged shared representations to improve task performance, while the Pareto-optimal solution was able to resolve conflicting knowledge between tasks. Additionally, we evaluated the generalizability of our framework through transfer learning by applying knowledge learned from the CRC dataset to other cancer datasets in miRTarBase [33]. This strategy overcame the challenges of limited data availability and expanded the application of miRNA target prediction to multiple cancer types. Our results demonstrated the effectiveness of multi-task and transfer learning in advancing miRNA research and improving our understanding of CRC and other malignancies. **Figure 1** presents the overall diagram of the proposed framework. The key contributions of this paper are summarized as follows:

**Figure 1.**
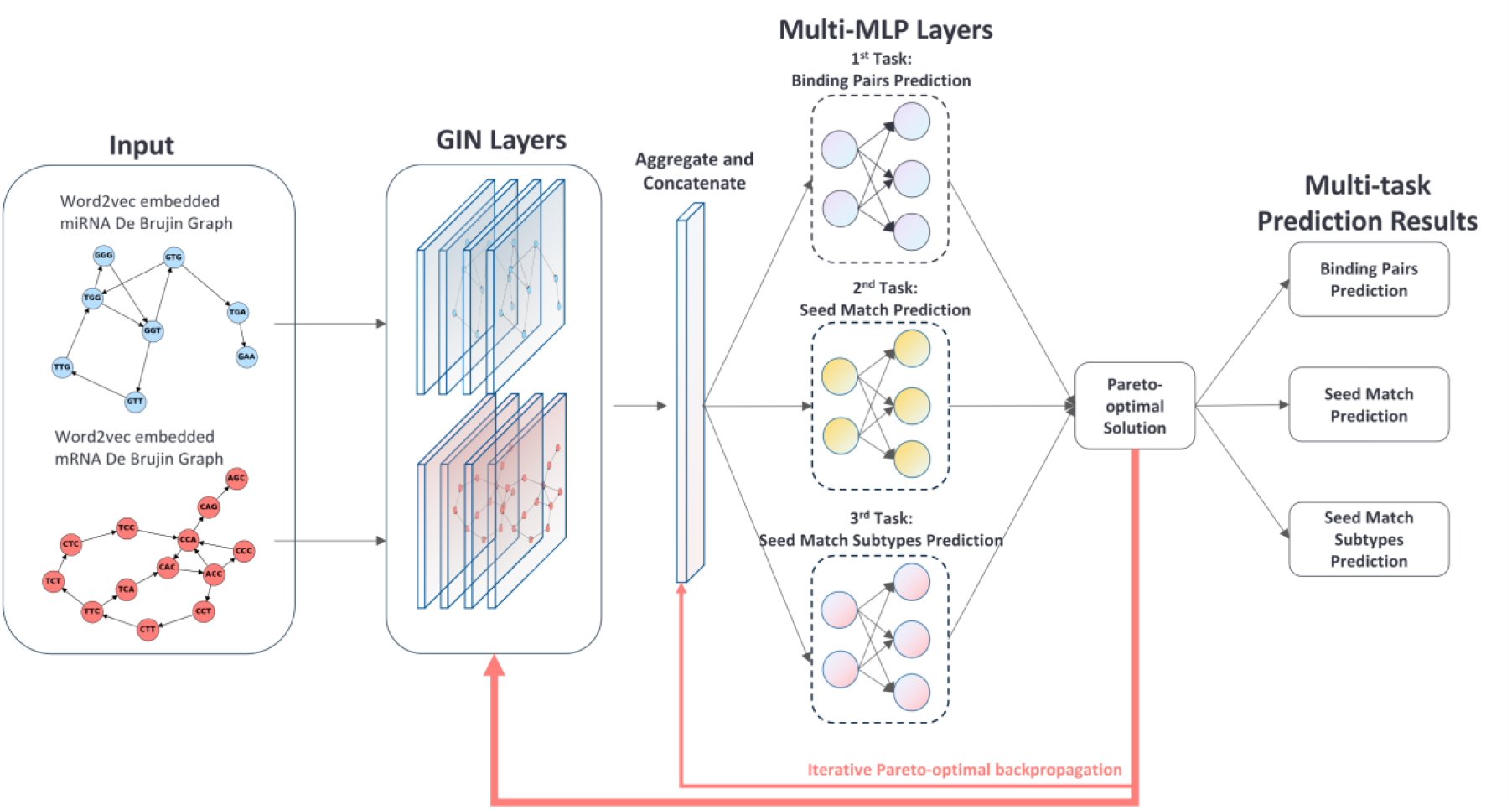
Graph-based Pareto-optimal multi-task learning framework for identifying miRNA-target interactions in colorectal cancer, consisting of three prediction tasks: (1) binding pairs prediction, (2) seed-match prediction, and (3) seed-match subtype prediction.

- We developed a unified multi-task learning framework that jointly predicts miRNA-mRNA binding pairs, seed match presence, and seed match subtypes, enabling both coarse and detailed regulatory inference for miRNA-target interactions in CRC.
- We introduced a graph-based Pareto-optimal solution strategy that dynamically mitigates task conflicts during training, effectively preventing any single objective from dominating the training process.
- We demonstrated the generalizability of the proposed framework by employing transfer learning to predict miRNA–target binding pairs interactions in other cancer types with limited data, achieving superior performance compared to models trained exclusively on the target datasets.

## Materials & Methods

### Problem Formulation

We addressed the hierarchical multi-task classification challenge inherent to RNA sequence analysis by our proposed graph-based Pareto-optimal multi-task learning framework. This framework jointly predicted three interrelated prediction tasks: (1) miRNA-mRNA binding pairs prediction (1^st^ Level), (2) miRNA-mRNA seed match classification (2^nd^ Level), and (3) miRNA-mRNA seed match subtype classification (3^rd^ Level). For each task, we first converted the RNA sequence into a de Bruijn graph constructed from overlapping *k*-mer embeddings. Then, a Graph Isomorphism Network (GIN) was employed as the base classifier to extract the structural features from each graph and generate the preliminary prediction results. Subsequently, our Pareto-optimal approach dynamically adjusted the relative importance among each task, ensuring the performance of each task reached optimality without sacrificing others. Finally, we incorporated transfer learning techniques to test the generalizability of the model and mitigated data scarcity for specific tasks.

### Datasets

In this study, we obtained experimentally verified miRNA-mRNA interaction pairs in colorectal cancer (CRC) using AGO-CLASH [34] data from HCT116 cells [35,36] (NCBI accession GSE164634). Adapter sequences were removed using Cutadapt [37] (version 3.4). After that, the trimmed paired-end FASTQ files were merged with PEAR software [38] (version 0.9.6). Each FASTQ file was then processed with Fastx_collapser from the FASTX-Toolkit (version 0.0.14) [39] to collapse duplicate reads into a single representative sequence. Unique Molecular Identifiers (UMIs) were subsequently trimmed from both the 5′ and 3′ ends using Cutadapt [37]. The cleaned FASTA files from the CLASH experiments were processed using the Hyb pipeline [34] to identify miRNA-mRNA hybrids. To improve specificity, only hybrids with a minimum interaction energy (ΔG) below −11.1 kcal/mol were retained as binding pairs labeled as “positive” [40]. For negative controls, the cleaned FASTA files were mapped to a human transcript database using Bowtie2 [41]. Gene abundance was calculated based on the total mapped reads, and the top 100 most abundant genes that were not identified among the positive pairs were selected. Reads corresponding to these genes were extracted from the SAM files. These reads were then randomly paired with miRNAs from the positive set to generate negative miRNA-mRNA hybrids. For the construction of pre-training models, miRNA and mRNA sequences were extracted from RNAcentral miRbase [42] and Ensembl [43], respectively. We extracted miRNA sequences from RNAcentral on the mammalian species, including humans, mice, and hamsters, etc. After duplicate removal, 16,253 unique miRNA sequences were retained, ranging from 15nt to 30 nt. Similarly, we collected mRNA sequences from the Ensembl database [43], with the same host species as miRNA. We ended up with 1,090,566 mRNA sequences after filtering duplicates, which would be used to construct the pre-training mRNA model. The average length of the selected mRNA sequences is 2609.9 nt, where the longest mRNA is 123,179 nt and the shortest is 35 nt.

### Data Preprocessing

We used k-mer method to encode RNA sequences, where k-mers were treated as analogous to words in sentences, enabling natural representation of nucleotide context. RNA sequences were segmented into overlapping *k*-mer subsequences using a sliding window of length *k*, generating *N*−*k*+1 subsequences from an RNA sequence of *N* nucleotides. We employed two pre-trained models [27] miRNA2Vec and mRNA2Vec to encode these k-mer subsequences into vectors, with *k* from 3 to 6 for optimal results. As shown in **Figure 2**, the transformed embeddings were then used to construct de Bruijn graphs, which efficiently represent the overlapping k-mer subsequences while preserving local sequence patterns. In the de Bruijn graph, each node corresponds to an embedded k-mer vector, and directed edges connect sequential *k*-mers. The edges are weighted by their normalized frequency, providing a detailed representation of RNA sequence structure and patterns.

**Figure 2.**
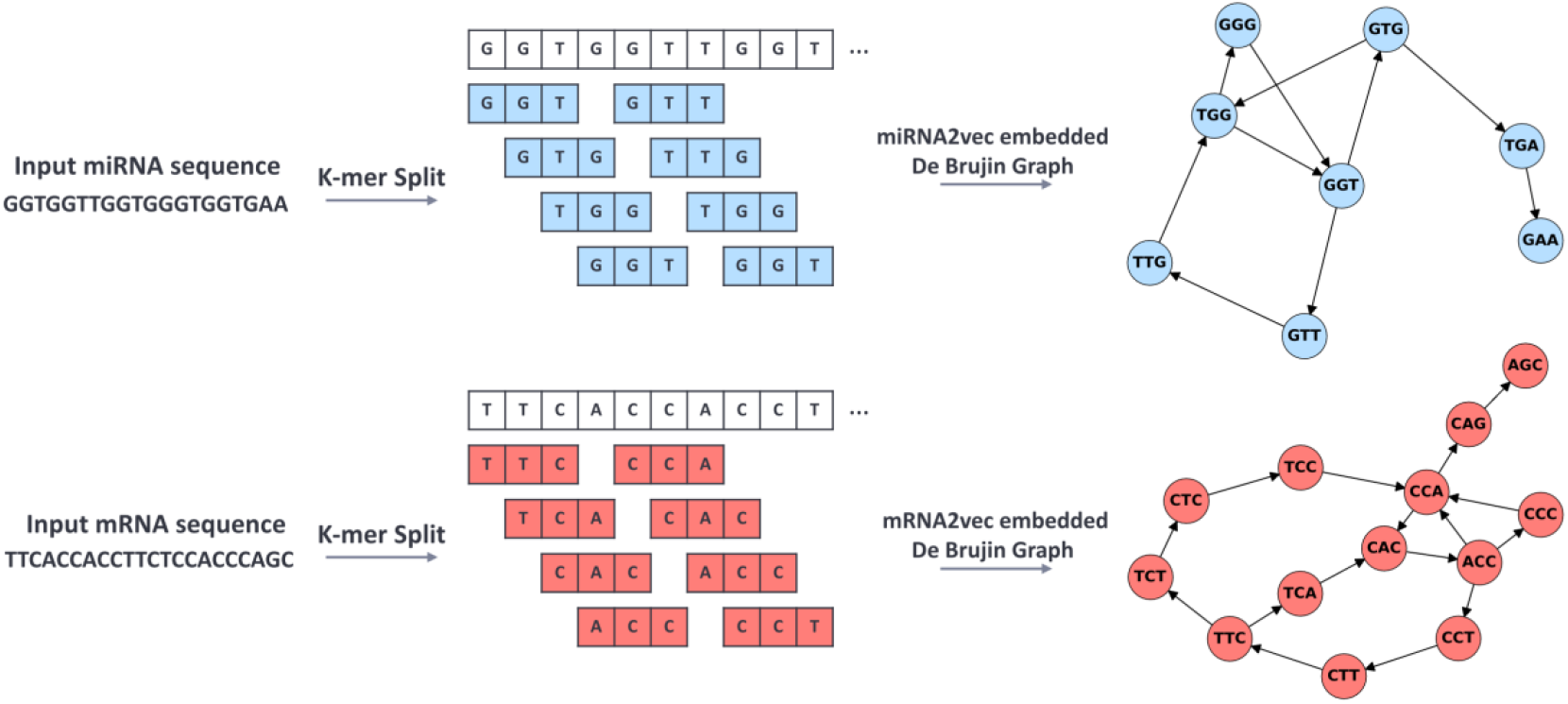
K-mer Generation and De Bruijn Graph Construction for miRNA and mRNA Sequences

### Graph Neural Networks (GNNs)

To effectively capture the RNA sequential information in the embedded de Bruijn graphs for each subtask, we employed graph neural networks (GNNs), which are well-suited for modeling complex graph structures. Building on our prior work [27], we selected graph isomorphism networks (GINs) [44] for the multi-task learning framework, as they demonstrated superior performance in miRNA-target binding pairs prediction compared to other GNN architectures, such as graph convolutional networks (GCNs) [45] and graph attention networks (GATs) [46]. GIN is designed to learn embeddings of graphs that are invariant under graph isomorphism [44] and it employs an iterative message passing and aggregation framework. At each iteration *k*, the hidden representation of a node *v*, denoted as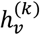, is updated by combining its current state with information aggregated from its neighbors:

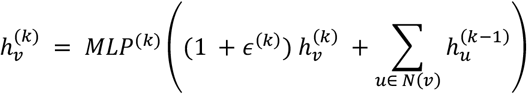

where *N*(*v*) represents the neighborhood of node *v* and *ε*^(*k*)^ is a parameter introduced to preserve permutation invariance. At iteration *k*, the node representation is computed by first aggregating its current representation with those of its neighbors. The multi-layer perceptron at iteration k, denoted as *MLP*^(*k*)^, then applies a nonlinear transformation to this aggregated sum and produces the updated node representation. This design enables the GIN to capture complex and nonlinear relationships within the local graph structure, ensuring that the resulting embeddings remain invariant for isomorphic graphs regardless of node ordering or labeling. While GINs effectively extract rich features from de Bruijn graphs, conflicts among gradients from different tasks can reduce the overall learning performance, motivating the need for a strategy that harmonizes the optimization process.

### Pareto-optimal multi-task framework

To address the hierarchical multi-task classification challenge inherent to miRNA-target interactions, we proposed a Pareto-optimal multi-task learning framework. This framework integrated the concepts of gradient-based optimization and multi-task learning to balance the trade-offs among the tasks, ensuring efficient resource allocation while optimizing the performance of each task. The model was jointly trained on three hierarchical classification levels: miRNA-mRNA binding pairs prediction (1^st^ Level), miRNA-mRNA seed match identification (2^nd^ Level), and seed match subtypes prediction (3^rd^ Level). At each iteration k, the proposed framework computed individual cross-entropy loss functions for each task, denoted as 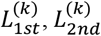, and 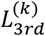. The total loss for the framework can be calculated as:

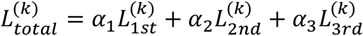

where the weights vector *α* = [*α*1, *α*2, *α*3]^*T*^(∑_*i*_ *α*_*i*_ = 1) is dynamically optimized using the Frank-Wolfe algorithm [47]. This adaptive weighting strategy ensures that, during each training epoch, no single task dominates the optimization process and that conflicts among task gradients are effectively balanced. The Frank-Wolfe solver addresses this problem by iteratively minimizing the gradient balancing objective function, formulated as:

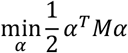

subject to:

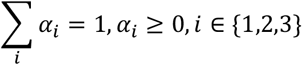

where *M* is the pairwise gradient similarity matrix, representing the inner products of gradients between all tasks denoted by:

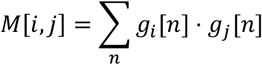

where *g*_*i*_ [*n*] and *g*_*j*_ [*n*] are the gradients of task *i* and *j*, respectively, for the *n*-th parameter of the model. The gradient similarity matrix *M*[*i, j*] quantifies the alignment between tasks: *M*[*i, j*] > 0 indicates that gradients of task *i* and *j* are aligned (point in similar directions), while *M*[*i, j*] < 0 indicates conflicting gradients. To solve the optimization problem, the Frank-Wolfe algorithm is employed to iteratively update the task weight *α*. At each iteration *t*, the update is performed as follows:

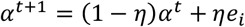

where *e*_*i*_ is a one-hot vector corresponding for task *i*, and *η* = 2/(*t* + 2) is the step size at iteration *t*. The Frank-Wolfe algorithm has a sublinear convergence rate 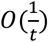, meaning that as the number of iterations increases, the improvement per iteration becomes smaller and most of the performance gains are achieved within the first few iterations. In practice, we observed that the first few iterations (e.g., 5~10) yield most of the performance gains, while additional iterations lead to diminishing returns. Therefore, in our proposed framework, we set *t* = 10 to balance optimization quality with computational efficiency. By updating the task weights *α* based on the gradient similarity matrix *M* in each training epoch *k*, our approach ensures that every gradient step approaches a Pareto-optimal solution, effectively harmonizing gradient conflicts across tasks and promoting balanced multi-task learning.

### Experimental Setup

#### Model Implementation and Evaluation

The models were implemented by PyTorch (v2.4.0) [48] and DGL (v2.4.0) [49]. The embeddings were generated with the Gensim Python package [50]. The dataset consists of paired miRNA-mRNA samples for three tasks: binding pairs prediction, seed match classification, and seed match subtypes classification. For the task of binding pairs prediction, interacting pairs were labeled as “1” (positive samples) and non-interacting pairs were labeled as “0” (negative samples). For the task of seed match prediction, seed match pairs were labeled as “1” (positive samples) and non-seed match pairs were labeled as “0” (negative samples). For the task of seed match subtypes prediction, numeric labels (0 to 4) were assigned to five distinct subtypes of seed matches: 8-mer, 7-mer-m8, 7-mer-A1, 6-mer, and offset-6-mer. The dataset was split into training (90%) and testing (10%) subsets using stratified sampling, ensuring that the proportion of each category is the same in both the training and testing sets. For our proposed model, we applied a minimum batch size of 128 for optimization. The learning rate is 0.001. All the models were trained for 300 epochs with an early stopping criterion of 60 epochs based on the weighted F1-score across all tasks. To prevent overfitting, a dropout rate of 0.3 was applied during training. Losses for each task were computed using task-specific cross-entropy loss functions, and the Frank-Wolfe algorithm dynamically optimized task weights to ensure Pareto-optimal learning across hierarchical levels. For evaluation, the testing set was used to assess the models using various metrics, including accuracy, precision, recall, F1-score, and the area under the Receiver Operating Characteristic curve (AUC). These metrics were computed using functions from the Scikit-learn [51] library to ensure consistency and robustness in performance evaluation.

#### Benchmark Comparison

To comprehensively assess our proposed GIN-CRC-Pareto framework, we designed multiple comparative benchmarks with ablation study, including (1) benchmarking against existing deep learning models, (2) internal comparisons with baseline variants of our method (single-task and two-task variants), and (3) ablation studies with fixed task weights that evaluate the performance difference between Pareto-optimal dynamic weights balancing and static weight configurations. First, we compared its performance with two categories of baseline methods. The first category is Classic Deep Learning Algorithms: Convolutional Neural Networks (CNNs), Recurrent Neural Networks with Gated Recurrent Unit (GRU), Combinations involving attention mechanisms: CNN + Attention, GRU + Attention. The second category is state-of-the-art miRNA target prediction models: preMLI [26], LncmirNet [52], Pmlipred [53], and PmliHFM [54]. For the baseline comparison models, we followed the original implementation settings provided in their respective publications and ensured that training and testing datasets were identical to those used for our framework. Second, we constructed and trained the following model variants under identical experimental conditions: the proposed three-task model (GIN-CRC-Pareto) and three two-task variants: GIN-CRC-Pareto-2-Task (1^st^ & 2^nd^ Tasks), GIN-CRC-Pareto-2-Task (1^st^ & 3^rd^ Tasks), GIN-CRC-Pareto-2-Task (2^nd^ & 3^rd^ Tasks), as well as the single-task models: independent GIN classifiers trained on each task. These models allowed us to assess how the inclusion or exclusion of specific tasks affects the model performance and generalization capability. Third, to further highlight the contribution of Pareto-optimal dynamic task balancing, we conducted ablation studies using fixed task weight settings. In the proposed GIN-CRC-Pareto framework, the loss function *L*_*total*_ is defined as:

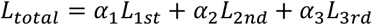

Here, *L*_1*st*_, *L*_2*nd*_, and *L*_3*rd*_ are calculated by the cross-entropy loss function through GIN classifier in each task, *α* = [*α*1, *α*2, *α*3]^T^(∑ _*i*_ *α* _*i*_ = 1) is the weight vector to balance the importance of each task. In our framework, the weight vector *α* was dynamically adjusted to follow the Pareto-optimal direction in each iteration. For the ablation study, we fixed the weight for the 2^nd^-Level task and systematically varied the weights of the 1^st^-Level and 3^rd^-Level tasks (*α*_1_ and *α*_3_) such that *α*_1_ + *α*_2_ + *α*_3_ = 1. In this study, *α*2 was fixed at 0.3 to allow flexibility in analyzing performance trends resulting from variations in *α*_1_ and *α*_3_. We set six combinations of *α*_1_ and *α*_3_, that is, {(0.1, 0.6), (0.2, 0.5), (0.3, 0.4), (0.4, 0.3), (0.5, 0.2), (0.6, 0.1)}, to examine how different fixed task weight allocations affect overall multi-task performance. This experiment enables a direct comparison between fixed weight allocation and dynamic Pareto-optimized weight adjustments, evaluating the robustness of our Pareto-optimized strategy in harmonizing conflicting gradients and promoting balanced performance across tasks.

#### Model Generalizability

To further test the generalizability of our proposed framework, we applied it to other cancers with limited miRNA-target interactions data. Specifically, we developed a transfer learning version of GIN-CRC-Pareto, **GIN-CRC-Transfer**, by fine-tuning our model that was pre-trained on CRC dataset using miRNA-target cell lines from cancers (e.g., bladder, breast, glioma, hepatocellular, lung, and prostate) collected from miRTarBase [33] that were experimentally validated. In typical GNN models, the initial convolutional layers extract macro-level information, such as shared structural or sequence features, while the subsequent layers, including the graph pooling layer and fully connected layers, specialize in task-specific knowledge relevant to classification tasks. Here, we utilized transfer learning by freezing the initial convolutional layers of the GIN, ensuring that the general shared knowledge from the CRC dataset was retained, such as common RNA sequential patterns. The subsequent layers of the GNN, which learn task-specific features, were fine-tuned to adapt to the cancer-specific information of the new datasets. This approach allowed the model to efficiently adapt to the new classification tasks while reducing the risk of overfitting, particularly when the model is adapted from a large dataset to small datasets. To generate negative data labels, we randomly sampled equal number of non-interacting miRNA-mRNA pairs from CRC HCT116 cells corresponding to each cancer type. We ended up with a total of 2,400 miRNA-mRNA pairs, involving 224 pairs from bladder cancer, 622 from breast cancer, 402 from glioma, 614 from hepatocellular carcinoma, 200 from lung cancer, and 338 from prostate cancer. For each cancer type, 80% of the data were used for training and the remaining 20% were reserved for evaluation.

In our experimental setup, learning rates, regularization coefficients, and other hyperparameters were recalibrated to optimize prediction performance. To prevent overfitting, particularly in the context of limited data, we employed early stopping and dropout strategies. All other model parameters were kept consistent with the original model. We evaluated three transfer learning configurations: 1-GIN-layer-freeze (only the first GIN layer frozen), All-GIN-layer-freeze (all GIN layers frozen), and No-freeze (no layers frozen). As the No-freeze setting consistently outperformed the others, we present its results as the representative configuration for GIN-CRC-Transfer. To further assess the transferability of the proposed framework, we introduced two baseline comparisons. The first, **No-Transfer**, involved training a GIN model from scratch on each target dataset, without utilizing prior knowledge, serving to highlight the benefit of transfer learning for small, domain-specific datasets. The second baseline, **GIN-CRC-Direct-Apply**, applied the pre-trained GIN-CRC-Pareto model pretrained on the CRC dataset directly to the target datasets without fine-tuning, allowing us to quantify the impact of the adaptation process. These comparisons ensure a comprehensive evaluation of the effectiveness and adaptability of our proposed transfer learning framework across diverse cancer types.

## Results

### Comparative performance between our model and deep learning benchmarks

We evaluated the performance of our proposed GIN-CRC-Pareto framework by benchmarking it against several widely used deep neural networks, including CNN, GRU, and their variants with attention mechanisms, as well as Transformer-based models. All models were trained and tested under identical experimental settings using the same pre-trained 5-mer embedded features, which have been identified as optimal for constructing graph-based representations of RNA sequences [27]. The results, summarized in

**Table 1**, showed that our proposed GIN-CRC-Pareto outperformed the competing models across multiple evaluation metrics. Specifically, for the 1^st^ task, our model achieved the highest accuracy (0.909), F1-score (0.909), precision (0.909), recall (0.909), and AUC (0.968). In comparison, the strongest baseline model, GRU, achieved an accuracy of 0.854 and AUC of 0.928, indicating a substantial performance gap. While the CNN model underperformed relative to the RNN model, adding attention did not yield noticeable improvements for CNN. This suggests that CNNs might lack the capacity to capture the sequential and relational complexity inherent in RNA interactions data, even with attention mechanisms. For the 2nd Task (miRNA-mRNA seed match prediction), GIN-CRC-Pareto again achieved the highest overall accuracy (0.853), F1-score (0.828), Precision (0.823), and AUC (0.924). Notably, both GRU and GRU+Attention achieved similar performance (accuracy = 0.851) compared to our model. Transformers performed significantly worse on this task (accuracy = 0.750, AUC = 0.815), likely due to the limited dataset size. In the more challenging 3rd Task (miRNA-mRNA seed match subtype prediction), GIN-CRC-Pareto maintained competitive performance (accuracy = 0.823, F1-score = 0.735), although GRU+Attention surpassed it in each metric (accuracy = 0.862, F1-score = 0.786, precision = 0.782, recall = 0.790). GRU-based models, especially with attention mechanisms, showed improved performance on this fine-grained classification task, indicating that RNN structures with attention might better capture subtle variations in seed pairing types. One potential reason for the slight drop in GIN-CRC-Pareto’s performance could be the limited number of samples within each subtype, making it harder for a graph-based model to capture the subtle differences between each subtype. Nevertheless, these results highlighted the robustness and effectiveness of our multi-task learning approach across different prediction objectives. Notably, the Transformer model underperformed across all metrics, particularly in the 2^nd^ and 3^rd^ tasks, suggesting that such architectures may require larger datasets to reach competitive performance. Overall, our proposed GIN-CRC-Pareto has demonstrated superior performance for multi-task miRNA target prediction, validating the utility of graph-based learning approaches in modeling complex RNA sequence relationships.

**Table 1.**
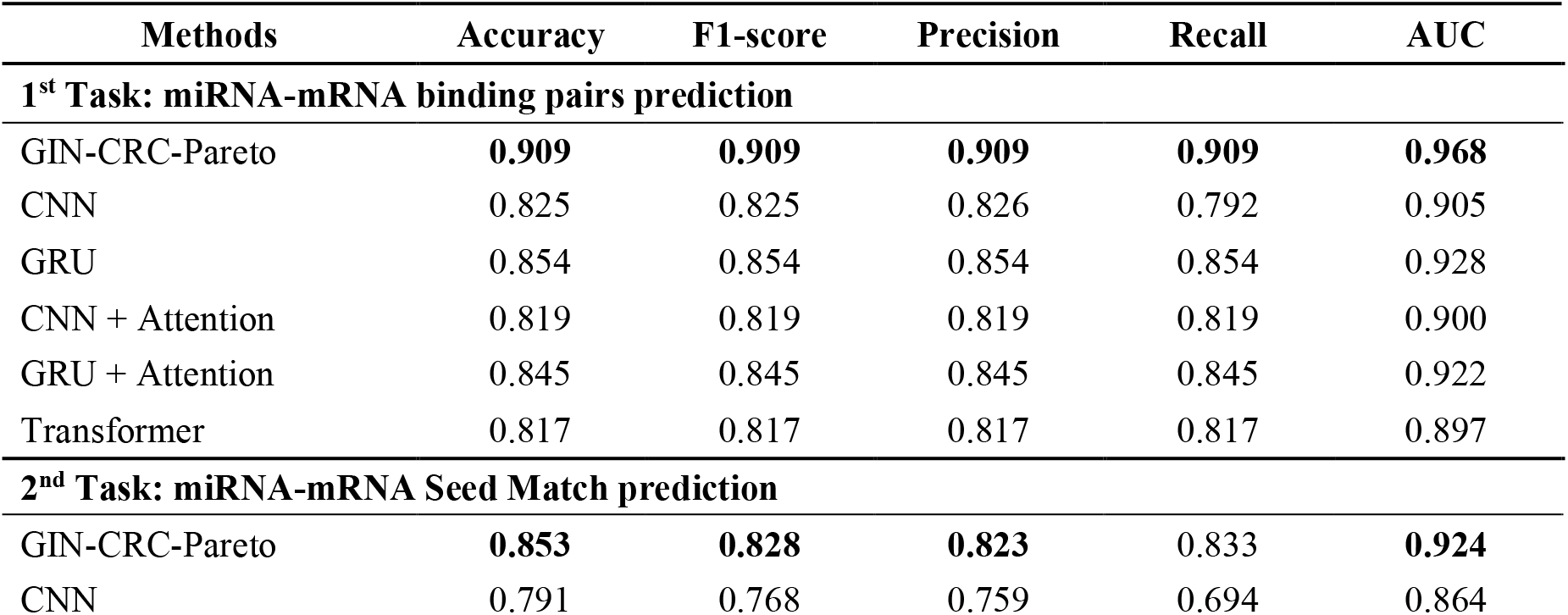
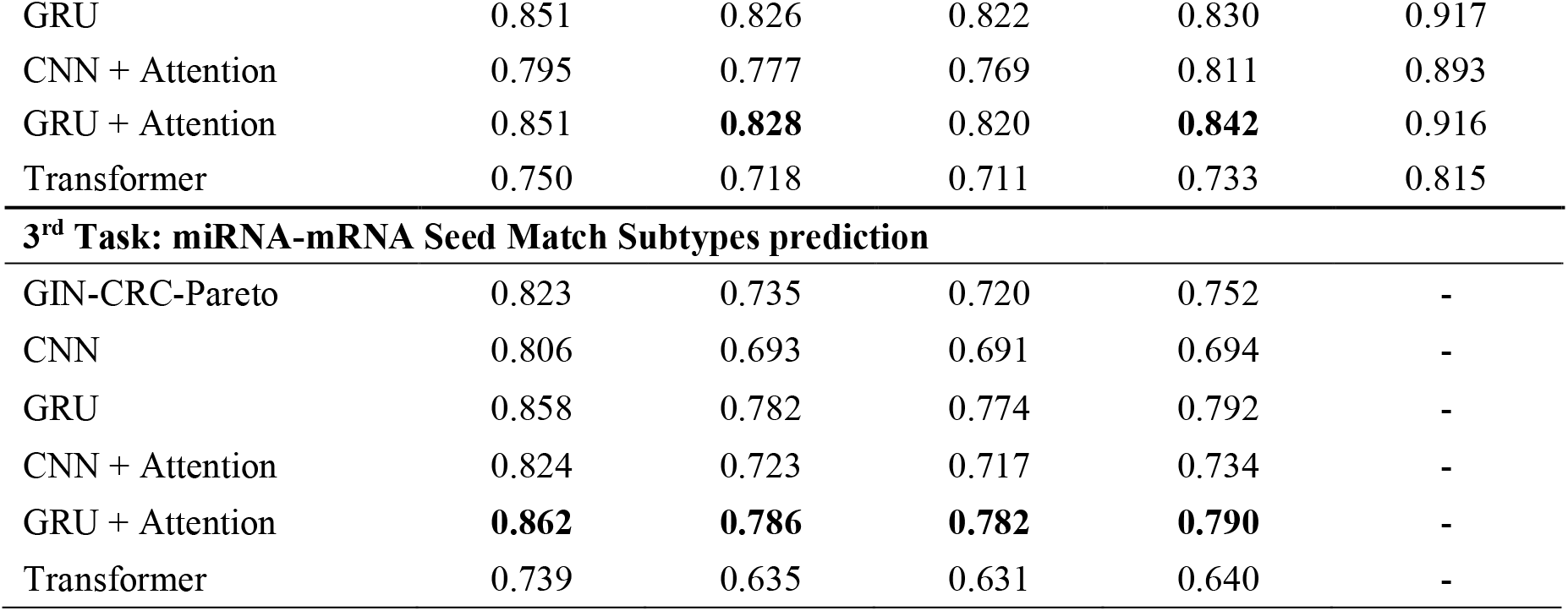
The performance comparison between our proposed framework with other state-of-the-art deep learning models on independent testing set. The AUC metric is not reported for the 3^rd^ Task as it is a multi-class problem. One-vs-All (OvA) AUCs for each subtype in 3^rd^ Task are detailed in Figure 3.

| Methods | Accuracy | F1-score | Precision | Recall | AUC |
| --- | --- | --- | --- | --- | --- |
| **1<sup>st</sup> Task: miRNA-mRNA binding pairs prediction** |  |  |  |  |  |
| GIN-CRC-Pareto | **0.909** | **0.909** | **0.909** | **0.909** | **0.968** |
| CNN | 0.825 | 0.825 | 0.826 | 0.792 | 0.905 |
| GRU | 0.854 | 0.854 | 0.854 | 0.854 | 0.928 |
| CNN + Attention | 0.819 | 0.819 | 0.819 | 0.819 | 0.900 |
| GRU + Attention | 0.845 | 0.845 | 0.845 | 0.845 | 0.922 |
| Transformer | 0.817 | 0.817 | 0.817 | 0.817 | 0.897 |
| **2<sup>nd</sup> Task: miRNA-mRNA Seed Match prediction** |  |  |  |  |  |
| GIN-CRC-Pareto | **0.853** | **0.828** | **0.823** | 0.833 | **0.924** |
| CNN | 0.791 | 0.768 | 0.759 | 0.694 | 0.864 |
| GRU | 0.851 | 0.826 | 0.822 | 0.830 | 0.917 |
| CNN + Attention | 0.795 | 0.777 | 0.769 | 0.811 | 0.893 |
| GRU + Attention | 0.851 | **0.828** | 0.820 | **0.842** | 0.916 |
| Transformer | 0.750 | 0.718 | 0.711 | 0.733 | 0.815 |
| **3<sup>rd</sup> Task: miRNA-mRNA Seed Match Subtypes prediction** |  |  |  |  |  |
| GIN-CRC-Pareto | 0.823 | 0.735 | 0.720 | 0.752 | - |
| CNN | 0.806 | 0.693 | 0.691 | 0.694 | - |
| GRU | 0.858 | 0.782 | 0.774 | 0.792 | - |
| CNN + Attention | 0.824 | 0.723 | 0.717 | 0.734 | - |
| GRU + Attention | **0.862** | **0.786** | **0.782** | **0.790** | - |
| Transformer | 0.739 | 0.635 | 0.631 | 0.640 | - |

### Comparative performance between multi- and single-task models

We compared the performance of our proposed model under different task configurations: a three-task configuration (GIN-CRC-Pareto), a two-task variant (GIN-CRC-Pareto-2-Task), and a single-task baseline (GIN-CRC-Single-Task). The Pareto-optimal solution is not applied to single-task setup as there are no conflicting problems to resolve. Each configuration was trained using the same pre-trained 5-mer embeddings and consistent experimental settings. The results in **Table 2** indicated the superiority of the multi-task framework compared to the single-task model. For the 1^st^ Task (miRNA-mRNA binding pairs prediction), the full GIN-CRC-Pareto achieved the best performance in all metrics (accuracy = 0.909, F1-score = 0.909, precision = 0.909, recall = 0.909, and AUC = 0.968). This indicated the multi-task framework’s ability to robustly capture complex interaction patterns between miRNAs and targets. For the 2^nd^ Task (seed match identification), the full GIN-CRC-Pareto model again outperformed others, with an accuracy of 0.853 and AUC of 0.924. Both two-task variants (1^st^ & 2^nd^ and 2^nd^ & 3^rd^) and the single-task model showed comparable but consistently lower performance. Interestingly, for the 3^rd^ Task, the two-task variant involving the 1^st^ and 2^nd^ Tasks exhibited lower performance than the single-task model, suggesting that incorporating information from the 2^nd^ Task may negatively impact the performance of the 3^rd^ Task. Conversely, the two-task variant involving the 1st and 3^rd^ Tasks (GIN-CRC-Pareto-2-Task (1^st^ & 3^rd^)) slightly outperformed the full three-task model on the 3^rd^ Task (accuracy = 0.831, precision = 0.747), indicating that excluding the 2^nd^ Task may allow the model to better focus on the 3^rd^ Task. However, this variant showed a slight decrease in the AUC of the 1st Task (0.962 vs. 0.968) compared to the three-task configuration, suggesting that the full three-task setting exhibited better generalization ability. Overall, the results highlighted the strength of the Pareto-optimal multi-task framework, which consistently matched or exceeded the performance of single-task approaches, underscoring the benefits of multi-task learning in miRNA target prediction problems.

**Table 2.**
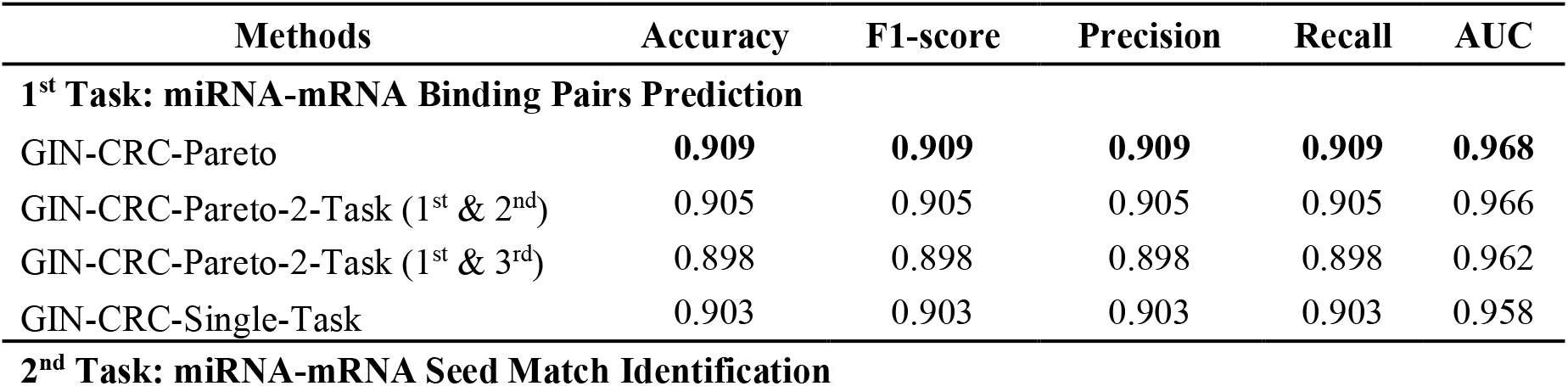
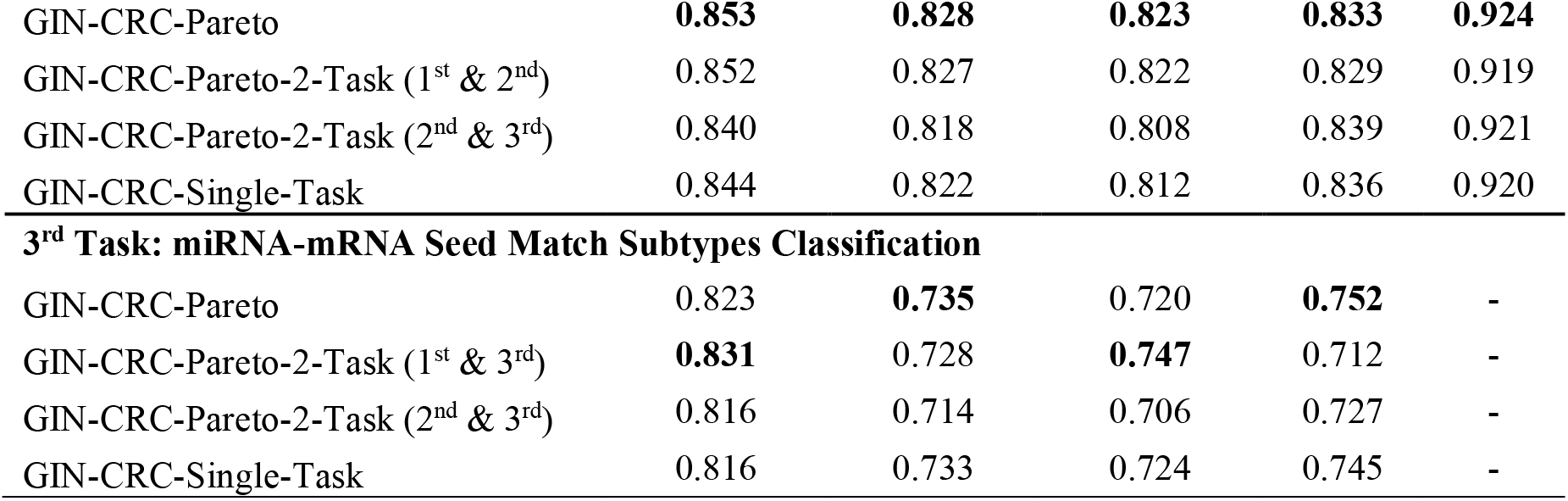
Comparative performance between the multi-task framework and the single-task model.

| Methods | Accuracy | F1-score | Precision | Recall | AUC |
| --- | --- | --- | --- | --- | --- |
| **1<sup>st</sup> Task: miRNA-mRNA Binding Pairs Prediction** |  |  |  |  |  |
| GIN-CRC-Pareto | **0.909** | **0.909** | **0.909** | **0.909** | **0.968** |
| GIN-CRC-Pareto-2-Task (1<sup>st</sup> & 2<sup>nd</sup>) | 0.905 | 0.905 | 0.905 | 0.905 | 0.966 |
| GIN-CRC-Pareto-2-Task (1<sup>st</sup> & 3<sup>rd</sup>) | 0.898 | 0.898 | 0.898 | 0.898 | 0.962 |
| GIN-CRC-Single-Task | 0.903 | 0.903 | 0.903 | 0.903 | 0.958 |
| **2<sup>nd</sup> Task: miRNA-mRNA Seed Match Identification** |  |  |  |  |  |
| GIN-CRC-Pareto | **0.853** | **0.828** | **0.823** | **0.833** | **0.924** |
| GIN-CRC-Pareto-2-Task (1<sup>st</sup> & 2<sup>nd</sup>) | 0.852 | 0.827 | 0.822 | 0.829 | 0.919 |
| GIN-CRC-Pareto-2-Task (2<sup>nd</sup> & 3<sup>rd</sup>) | 0.840 | 0.818 | 0.808 | 0.839 | 0.921 |
| GIN-CRC-Single-Task | 0.844 | 0.822 | 0.812 | 0.836 | 0.920 |
| **3<sup>rd</sup> Task: miRNA-mRNA Seed Match Subtypes Classification** |  |  |  |  |  |
| GIN-CRC-Pareto | 0.823 | **0.735** | 0.720 | **0.752** | - |
| GIN-CRC-Pareto-2-Task (1<sup>st</sup> & 3<sup>rd</sup>) | **0.831** | 0.728 | **0.747** | 0.712 | - |
| GIN-CRC-Pareto-2-Task (2<sup>nd</sup> & 3<sup>rd</sup>) | 0.816 | 0.714 | 0.706 | 0.727 | - |
| GIN-CRC-Single-Task | 0.816 | 0.733 | 0.724 | 0.745 | - |

### Comparison between GIN-CRC-Pareto and existing tools

We further benchmarked our proposed framework against several existing state-of-the-art methods across all three tasks. As most public models primarily focus on miRNA–target binding pairs prediction (1^st^ Task), with limited support for 2^nd^ Task and 3^rd^ Task, we adapted these models to enable comparative evaluation across the full task spectrum. The comparative models included preMLI [26], LncmirNet [52], Pmlipred [53], PmliHFM [54], and Gra-CRC-miRTar [27]. As illustrated in **Figure 3**, the proposed GIN-CRC-Pareto framework demonstrates consistent and superior performance across all three levels of miRNA-target interaction prediction tasks. At the 1^st^-Level Task, GIN-CRC-Pareto achieved an AUC of 0.968, matching the performance of Gra-CRC-miRTar model and slightly outperforming all other baseline models. In the 2^nd^-Level Task, GIN-CRC-Pareto maintained high performance with an AUC of 0.924. All models showed reasonably strong performance. PmliHFM reached the highest AUC of 0.942 in the 2^nd^-Level Task and slightly outperformed GIN-CRC-Pareto, but the performance of PmliHFM dropped considerably in the 1^st^-Level Task compared to Gra-CRC-miRTar (AUC = 0.899 vs 0.968), suggesting the limited generalizability of PmliHFM across tasks. In the 3^rd^-Level Task, we compared the AUC of One-vs-All (OVA) classification task for five distinct seed match categories: 6-mer, 7-mer-A1, 7-mer-m8, 8-mer, and offset-6-mer. GIN-CRC-Pareto consistently outperformed other tools in nearly all categories. GIN-CRC-Pareto achieved strong performance with an AUC of 0.925, equivalent to Gra-CRC-miRTar (AUC = 0.925), while the remaining models exhibited moderate results in AUC, including LncMirNet (0.787), PmliPred (0.863), PmliHFM (0.837), and preMLI (0.782). Across other seed match subtypes, GIN-CRC-Pareto consistently obtained competitive AUCs ≥ 0.920, while other models showed performance declines (e.g., preMLI reached an AUC of 0.869 for 7-mer-m8 and LncMirNet scored 0.85 for offset-6-mer). These results highlight the robustness and adaptability of the GIN-CRC-Pareto framework. While some existing models achieved comparable results in certain individual tasks, their generalization ability across tasks was limited. In contrast, our multi-task learning approach consistently delivered high predictive performance across all hierarchical levels, demonstrating its effectiveness in capturing both global and context-specific features of miRNA target interactions.

**Figure 3.**
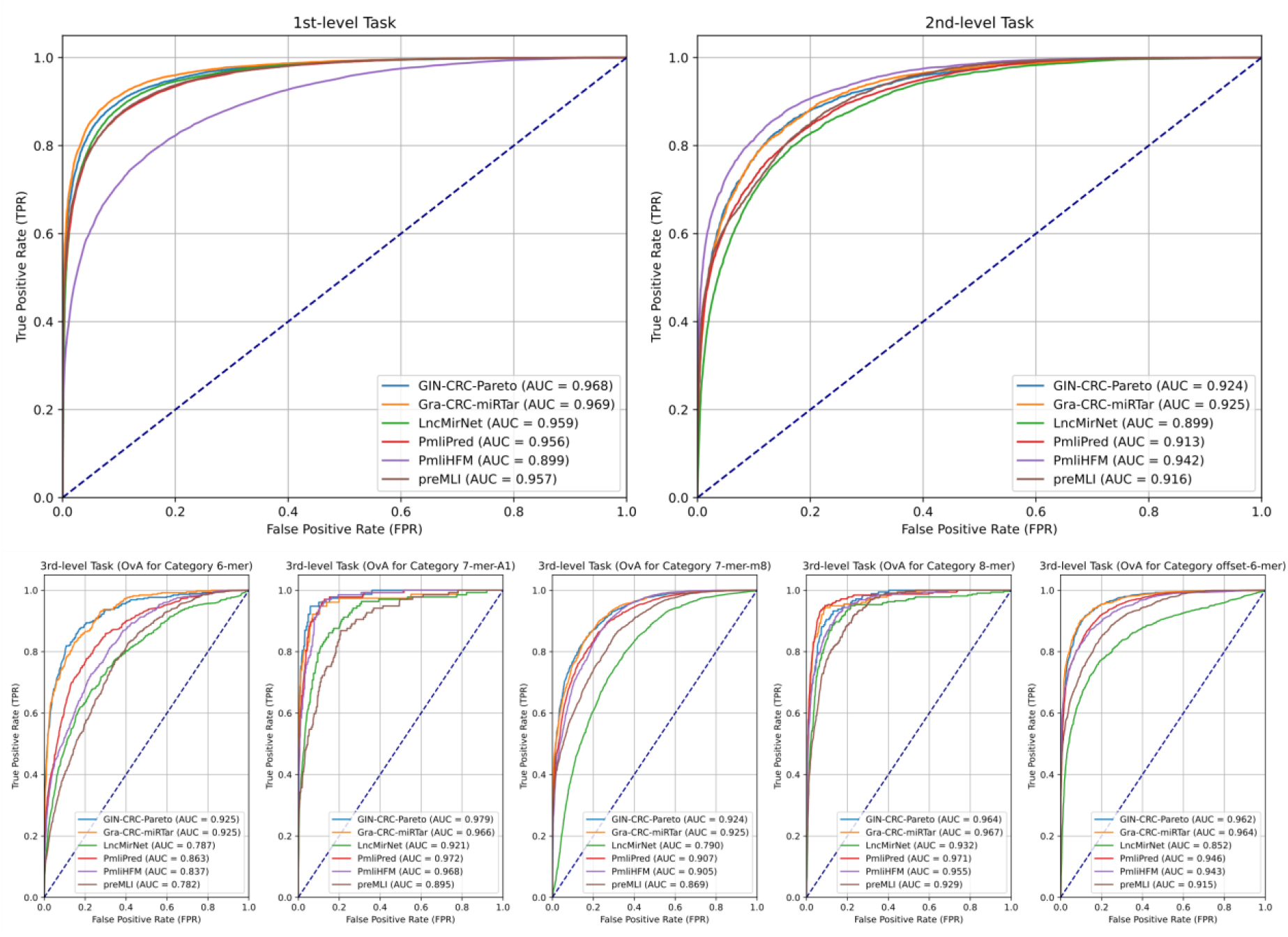
Comparative performance between our proposed framework and other existing tools on multi-task miRNA-target interaction identification.

### Ablation study between task weights tuning and Pareto-optimal solution

As shown in Table 3, we compared the performance between the Pareto solution and fixed weights of our multi-task learning (GIN-CRC-FW) for identifying miRNA-target prediction using our proposed framework. According to Table 3, GIN-CRC-Pareto consistently achieved the highest or comparable performance, demonstrating its effectiveness in optimizing multi-task objectives without compromising individual task performance. In more detail, for the 1^st^-Level Task, the accuracy and F1-score steadily increased as *α*_1_ increased from 0.1 to 0.6, reaching a peak of 0.909 at *α*_1_ = 0.6. However, this came at the expense of reduced performance in the 3^rd^-Level Task, where accuracy and F1-score dropped from 0.747 and 0.684. In contrast, the GIN-CRC-Pareto achieved the highest overall accuracy (0.909) and F1-score (0.909) for the 1st-Level Task, while also maintaining competitive performance in both the 2^nd^-Level Task (accuracy = 0.838, F1 = 0.818) and the 3^rd^-Level Task (accuracy = 0.823, F1 = 0.735). This is probably due to the dynamic optimization of the weights across multiple tasks, which adaptively balances trade-offs among tasks based on task conflicts and interdependencies. Notably, none of the fixed-weight configurations were able to simultaneously match or surpass the performance of the Pareto-optimal model across all tasks. These results highlighted the advantage of the Pareto-optimal approach in achieving robust and balanced predictions in multi-task learning scenarios without sacrificing individual task performance. The complete results involving other evaluation metrics, such as precision, recall, and AUC, were provided in **Table S1** of the Supplementary Material.

**Table 3.** Comparison study between GIN-CRC-Pareto and GIN-CRC-FW variants trained with fixed task weight configurations.

| Weights Comparison (fix $\alpha_2 = 0.3$) | Accuracy | | | F1-score | | |
| --- | --- | --- | --- | --- | --- | --- |
|  | 1<sup>st</sup> Task | 2<sup>nd</sup> Task | 3<sup>rd</sup> Task | 1<sup>st</sup> Task | 2<sup>nd</sup> Task | 3<sup>rd</sup> Task |
| GIN-CRC-FW ($\alpha_1 = 0.1, \alpha_3 = 0.6$) | 0.865 | 0.816 | 0.832 | 0.865 | 0.798 | 0.747 |
| GIN-CRC-FW ($\alpha_1 = 0.2, \alpha_3 = 0.5$) | 0.888 | 0.838 | 0.831 | 0.888 | 0.817 | 0.735 |
| GIN-CRC-FW ($\alpha_1 = 0.3, \alpha_3 = 0.4$) | 0.898 | 0.842 | 0.834 | 0.898 | 0.819 | 0.722 |
| GIN-CRC-FW ($\alpha_1 = 0.4, \alpha_3 = 0.3$) | 0.895 | 0.843 | 0.828 | 0.895 | 0.822 | 0.725 |
| GIN-CRC-FW ($\alpha_1 = 0.5, \alpha_3 = 0.2$) | 0.902 | 0.827 | 0.821 | 0.902 | 0.808 | 0.729 |
| GIN-CRC-FW ($\alpha_1 = 0.6, \alpha_3 = 0.1$) | 0.907 | 0.843 | 0.793 | 0.907 | 0.820 | 0.684 |
| GIN-CRC-Pareto (adaptive weights) | 0.909 | 0.838 | 0.823 | 0.909 | 0.818 | 0.735 |

### Performance Assessment across Other Cancer Types

To evaluate the generalizability of the proposed GIN-CRC-Pareto framework, we applied it to identify miRNA-target interaction leveraging transfer learning techniques across other six cancers, including bladder, breast, glioma, hepatocellular, lung, and prostatic. As shown in **Figure 4**, the performance of the GIN-CRC-Transfer approach was benchmarked against two baselines: No-Transfer (a GIN model trained from scratch on each target dataset) and GIN-CRC-Direct-Apply (the CRC-pretrained GIN-CRC-Pareto model applied directly without transfer learning). Across all target datasets, GIN-CRC-Transfer consistently outperformed both baselines in terms of accuracy and AUC, confirming the benefits of transfer learning in low-resource scenarios. In Lung cancer dataset, GIN-CRC-Transfer achieved the highest accuracy (0.895) and AUC (0.916), compared to No-Transfer (accuracy = 0.874, AUC = 0.888) and GIN-CRC-Direct-Apply (accuracy = 0.789, AUC = 0.860). On Hepatocellular dataset, it reached an accuracy = 0.928 and AUC = 0.955, significantly surpassing both baselines. In Glioma and Breast datasets, GIN-CRC-Transfer achieved accuracy scores of 0.918 and 0.925, respectively, with both AUC values exceeding 0.95, demonstrating strong adaptability to distinct cancer types. Overall, our proposed framework suggests significant generalizability capacity, incorporating transfer learning techniques to identify miRNA-target interaction pairs in different cancer types, making it a promising strategy for cross-cancer miRNA-target interaction prediction tasks.

**Figure 4.**
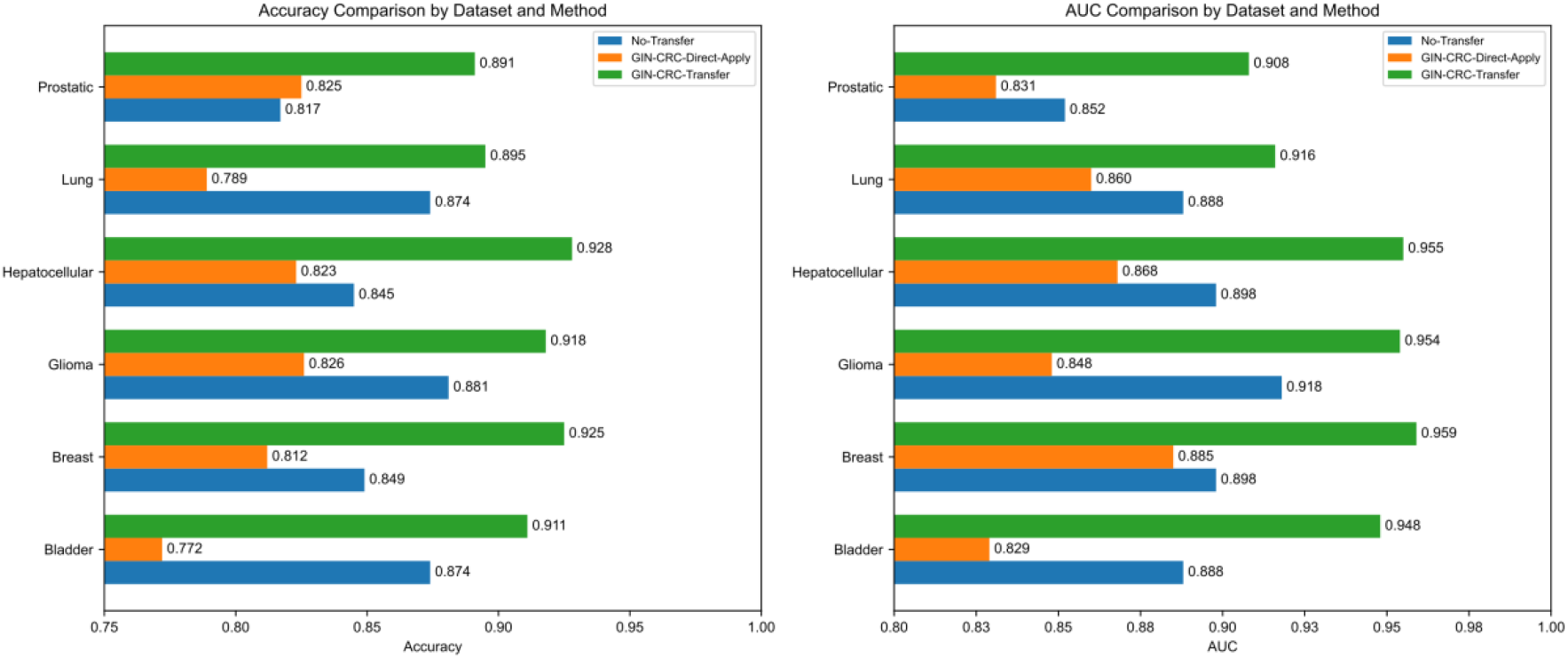
The generalizability assessment of our proposed framework to predict miRNA-target interaction across multiple cancer types.

## Discussion

The proposed GIN-CRC-Pareto framework demonstrated robust capability in miRNA-mRNA interactions prediction by effectively addressing the complexities inherent in multi-level tasks. By balancing miRNA-mRNA binding pairs prediction (1^st^-Level), seed match type identification (2^nd^-Level), and seed match subtype classification (3^rd^-Level) within a unified framework, the method achieved high performance without sacrificing individual task accuracy. Specifically, the framework attained accuracies of 0.909, 0.853, and 0.823 for the 1^st^, 2^nd^, and 3^rd^ level tasks, respectively. The accuracy decreased progressively from the 1st to the 3rd task due to reduced training data availability and potential error propagation in generating seed match and subtype labels. Despite this, GIN-CRC-Pareto consistently outperformed both conventional deep learning models and current state-of-the-art methods in miRNA–target interaction prediction. Though some baseline models perform well on individual tasks, they struggled to maintain balanced performance across multiple tasks. For instance, LncMirnet reached reasonable performance in the 1^st^ and 2^nd^ level tasks, but its performance dropped substantially on the 3^rd^ level task, with OvA AUCs of only 0.787 for the 6-mer category and 0.790 for the 7-mer-m8 category. We also compared single-task and two-task variant models with our proposed framework. The results indicate that single-task and two-task variant models achieved a comparably high performance on the 1^st^ task, likely due to the large dataset, which can provide efficient knowledge for effective learning. However, the performance of the variant models dropped notably on the 2^nd^ and 3^rd^ level tasks, where smaller datasets limited the models’ ability to fully capture task-specific patterns. In contrast, our proposed framework remained robust across all tasks. By leveraging shared information from the first task, it effectively supported learning in subsequent tasks, demonstrating its robustness and adaptability in addressing the varying demands of hierarchical classification levels. Although the 1^st^ task involves distinct labels and objectives, the gradient directions of its loss function often align with those of the 2^nd^ and 3^rd^ tasks during the early training stages, thereby enhancing their performance. This highlights the advantage of incorporating the Pareto-optimal approach, particularly in seed match subtype classification, where biologically important but underrepresented categories pose a challenge.

Interestingly, one of the two-task variant models, GIN-CRC-Pareto-2-Task (1st & 3rd), slightly outperformed the proposed GIN-CRC-Pareto on the 3^rd^ level task. It is suggested that excluding the 2^nd^ level task may help the model to concentrate better on the subsequent 3^rd^ level task. However, this variant exhibited a slight decline in the AUC of the 1st task (0.962 vs. 0.968), indicating that the complete three-task configuration provides better overall generalization by effectively leveraging the interdependencies among tasks. The benefits of this strategy were further confirmed through comparative studies of different weight tuning methods. When we fixed the weights allocation for the 2^nd^ level tasks and manually adjusted the weights for the 1^st^ and 3^rd^ level tasks, we observed that the accuracy and F1-score of individual tasks varied depending on the assigned weights. While manual adjustment of task weights could improve performance for a specific task, it often came at the expense of others. In contrast, the Pareto-optimal strategy consistently delivered balanced and high performance across all tasks by dynamically adjusting weights and effectively mitigating gradient conflicts. Moreover, the generalizability of the GIN-CRC-Pareto framework was further demonstrated to predict miRNA-target interactions across different cancer types using transfer learning techniques. By transferring knowledge learned from CRC, in which we have sufficient data for training the model, to a range of other cancer types within the miRTarBase repository, the proposed transfer learning framework GIN-CRC-Transfer achieved notable performance gains compared to both baselines: **GIN-CRC-Direct-Apply** and **No-Transfer**. Notably, even in the bladder dataset, where the training set was extremely limited (only 179 samples), GIN-CRC-Transfer attained a superior AUC of 0.948, significantly outperforming GIN-CRC-Direct-Apply (0.849) and No-Transfer (0.918). These results highlight the potential of the proposed framework’s ability to retain and adapt functional miRNA-mRNA binding patterns across different malignancies, even in scenarios with limited training data.

Despite its strengths, the proposed framework presents several limitations. One notable challenge lies in the added complexity of the Pareto-optimal approach, which introduces a more intricate model optimization process and increases the burden of hyperparameter tuning due to the need to balance trade-offs among multiple tasks. This complexity can lead to elevated computational costs and extended training times, particularly when dealing with a large hyperparameter search space or high-dimensional datasets. Another potential limitation is the current focus on a single species, i.e., humans, and limited incorporation of broader biological or clinical contexts. Expanding the framework to support cross-species prediction would enhance its applicability in translational and comparative genomics. Moreover, integrating multi-modal data, such as clinical features, gene expression profiles, or mutation status, could improve the biological interpretability and predictive accuracy of the model in disease-specific applications. Future work would explore strategies to mitigate these issues, such as incorporating more efficient optimization techniques, automated hyperparameter tuning methods (e.g., Bayesian optimization), or leveraging parallel and distributed computing infrastructures to accelerate training. These extensions would enable GIN-CRC-Pareto to serve as a more comprehensive tool for precision medicine, particularly in cancer biology and miRNA-based therapeutic discovery.

## Conclusion

In this study, we introduced GIN-CRC-Pareto, a graph-based Pareto-optimal multi-task learning framework designed to address the complex challenge of hierarchical miRNA-mRNA interaction prediction tasks involving binding pairs prediction, seed match type classification, and seed match subtype identification. By incorporating a Pareto-optimal gradient balancing strategy, the framework dynamically adjusts task weights during training to resolve conflicts between objectives and ensure balanced optimization across tasks. Experimental results demonstrated that GIN-CRC-Pareto achieves consistently decent performance across all three predictive tasks, outperforming traditional models that often struggle with task trade-offs. Additionally, we further validated the generalizability and scalability of the proposed framework by applying it for miRNA-target interaction prediction across multiple cancer types in combination with transfer learning techniques. Overall, GIN-CRC-Pareto offers a robust, generalized, and comprehensive solution for multi-level miRNA-mRNA interactions prediction tasks.

## ACKNOWLEDGMENTS

This work was supported by the Elsa U. Pardee Foundation; the University of Florida Health Cancer Center Pilot Grant [AGR DTD 01-30-2015]; the Florida Department of Health William G. “Bill” Bankhead, Jr., and David Coley Cancer Research Program; the National Institute of General Medical Sciences [R35GM128753]; and the National Science Foundation [CNS-2318210]. Daily maintenance and user management were strongly supported by staff members at the Research Computing Group at the University of Memphis. [55].

