## Supplementary for "GIN-CRC-Pareto: A graph-based Pareto-optimal multi-task learning framework to identify miRNA-target interactions in colorectal cancer"

**Table S1** Comparison study between task weights tuning and Pareto-optimal solution

| **Weights**  **Comparison (fix** $\boldsymbol{\alpha}_{\boldsymbol{2}}$ **= 0.3)** | **Accuracy** | | | **F1-score** | | | **Precision** | | | **Recall** | | | **AUC** | |
| --- | --- | --- | --- | --- | --- | --- | --- | --- | --- | --- | --- | --- | --- | --- |
|  | Task 1 | Task 2 | Task 3 | Task 1 | Task 2 | Task 3 | Task 1 | Task 2 | Task 3 | Task 1 | Task 2 | Task 3 | Task 1 | Task 2 |
| **GIN-3-task (**$\boldsymbol{\alpha}_{\boldsymbol{1}}$ **= 0.1,**  $\boldsymbol{\alpha}_{\boldsymbol{3}}$ **= 0.6)** | 0.865 | 0.816 | 0.832 | 0.865 | 0.798 | 0.747 | 0.865 | 0.787 | 0.758 | 0.865 | 0.827 | 0.744 | 0.941 | 0.914 |
| **GIN-3-task (**$\boldsymbol{\alpha}_{\boldsymbol{1}}$ **= 0.2,**  $\boldsymbol{\alpha}_{\boldsymbol{3}}$ **= 0.5)** | 0.888 | 0.838 | 0.831 | 0.888 | 0.817 | 0.735 | 0.888 | 0.806 | 0.733 | 0.888 | 0.834 | 0.740 | 0.957 | 0.919 |
| **GIN-3-task (**$\boldsymbol{\alpha}_{\boldsymbol{1}}$ **= 0.3,**  $\boldsymbol{\alpha}_{\boldsymbol{3}}$ **= 0.4)** | 0.898 | 0.842 | 0.834 | 0.898 | 0.819 | 0.722 | 0.898 | 0.810 | 0.732 | 0.898 | 0.830 | 0.727 | 0.962 | 0.916 |
| **GIN-3-task (**$\boldsymbol{\alpha}_{\boldsymbol{1}}$ **= 0.4,** $\boldsymbol{\alpha}_{\boldsymbol{3}}$ **= 0.3)** | 0.895 | 0.843 | 0.828 | 0.895 | 0.822 | 0.725 | 0.895 | 0.811 | 0.728 | 0.895 | 0.839 | 0.722 | 0.961 | 0.923 |
| **GIN-3-task (**$\boldsymbol{\alpha}_{\boldsymbol{1}}$ **= 0.5,** $\boldsymbol{\alpha}_{\boldsymbol{3}}$ **= 0.2)** | 0.902 | 0.827 | 0.821 | 0.902 | 0.808 | 0.729 | 0.902 | 0.797 | 0.734 | 0.902 | 0.835 | 0.727 | 0.964 | 0.921 |
| **GIN-3-task (**$\boldsymbol{\alpha}_{\boldsymbol{1}}$ **= 0.6,**  $\boldsymbol{\alpha}_{\boldsymbol{3}}$ **= 0.1)** | 0.907 | 0.843 | 0.793 | 0.907 | 0.820 | 0.684 | 0.908 | 0.811 | 0.675 | 0.907 | 0.831 | 0.698 | 0.967 | 0.919 |
| **GIN-3-task-Pareto** | 0.909 | 0.838 | 0.823 | 0.909 | 0.818 | 0.735 | 0.909 | 0.807 | 0.720 | 0.909 | 0.839 | 0.752 | 0.969 | 0.924 |
